# PERFORMANCE OF FOUR NEWLY RELEASED CASSAVA VARIETIES IN A FALLOWED LAND OVER TWO FARMING SEASONS

**DOI:** 10.1101/2025.01.01.631026

**Authors:** C. O. Ossai, S. A. Ojobor, T. O. Ogwuche, S. C. Akpeji, I. S. Alama, E. M. Diebiru-Ojo

## Abstract

Cassava (*Manihot esculenta* Crantz) is an important crop for humans due to its staple and industrial values. Currently the demand is more than the supply as the current output falls below the expected. This necessitated the breeding of superior genotypes that are high yielding. These superior genotypes also requires a fertile soil for optimum production. However, this has been primarily achieved through the application of inorganic manures. It is then important to evaluate four newly released cassava varieties in an organically enriched soil relative to the local best variety. Four newly released cassava varieties, Hope, Obasanjo2, Baba 70, and Game changer, and one Local Best (LB) was planted in 5-years fallowed farm lands at three plots (A, B and C) for 2 seasons. It was a three-way factorial (5-varieties*3-farms*2-seasons) arranged in a randomized complete block design and replicated three times. Data were taken on the Fresh Tuber Weight (FTW) and Stem Height (SH) per variety, and pre and post soil status. Data were analyzed using analysis of variance, while differences in varietal means were separated using least significant differences at 5% level of significance. The FTW and SH differed significantly across varieties, years and interaction between variety, farms and year, and ranged from 19.2±0.5 (LB) to 41.0±0.5 (Obasanjo2), 32.1±0.4 (farm C) to 34.1±0.4 (farm A), 30.4±0.3 (year two) to 35.0±0.3 (year one), and the interactions between variety and year, farms and year, and variety, year and farms were significant. The soil macro and micro elements declined in the post relative to the pre status. The yield obtained from the improved varieties almost doubled the local best and yield declined in the second season.

## INTRODUCTION

Cassava (*Manihot esculenta* Crantz) is an important staple for humans worldwide as its root tubers can be processed into different consumable forms like garri, eba, tapioca etc (1). Also, in some eastern part of Africa, the leaves of some cassava varieties serve as soup ingredients for consumption. Apart from the direct consumption of cassava products by humans, it has found its usefulness in various industries like the making of animal feeds and the production of biofuels (2). Due to the valuable uses of cassava, the demand for its products increased significantly from 251.5 million tons in 2010 to 302.7 million tons in 2020 (3). The major cassava producing countries in Sub-Saharan Africa (SSA) includes Nigeria, Democratic Republic of Congo (DRC), Angola, Ghana and Mozambique listed in decreasing order of output (4). In Nigeria, cassava is considered the most important food and industrial crop as it generates income for over 30 million Nigerian who engages in the cultivation, processing and the trading (5; 4).

The increased demand for cassava and its products in Nigeria ultimately leads to increased interest in the cultivation of high yielding cassava varieties in Nigeria and other parts of the world. However, cassava just like other food crops is dependent on the genotypic, environmental and genotype by environment interaction to combat the various biotic and abiotic factors militating against optimum yield in an area (6; 7). Despite Nigeria leading the world in the total cassava production, the current output in the country is far below its demand for food, raw material for industrial uses and export (8). This has necessitated agricultural research institutes like the International Institute of Tropical Agriculture (IITA) and the National Root Crop Research Institute (NRCRI) to constantly breed and develop several elite improved cassava varieties that can combat various biotic and abiotic stresses in order to improve their productivity and meet the teaming consumer demand (9). One of the major constraint to the full adoption of the improved cassava varieties in Nigeria is the awareness of the local farmers to the superior characteristics of the improved cassava varieties to the existing local varieties that has been cultivated for years despite reports showing that farmers who adopted improved cassava varieties had higher tuber yield relative to their old cultivated varieties (5). In most instances, local farmers had to wait and see the outcome of the produce from the early adopters to have full conviction to adopt the improved varieties (10).

While looking at the development and adoption of improved cassava varieties with sight on improved yield and productivity, it is important to consider the cultivating soil nutrient status as inappropriate cultivation of the soil can alter the genetic gain of the improved varieties. Supplementation of the soil nutrient content with inorganic fertilizer has been advocated to enhance cassava productivity (11; 12), but the indiscriminate application of the inorganic fertilizers can also result to severe environmental degradations (13). Hence, during the awareness campaign on the adoption of improved cassava, efforts should also be made to educate the local farmers on the appropriate timing and quantity of synthetic fertilizer to augment their soil nutrient status for efficient productivity (14). However, some farming communities like Ogbagu Ogume in Delta State practices organic nutrient replenishing system whereby the replenishment of their soil nutrient status is based on the practice of shifting cultivation with a minimum of 5 years before returning to a farmland that can be cultivated for a maximum of two seasons. Being a forest zone, within the 5 years of absence, plant leaves and debris are expected to have fallen and decayed in the uncultivated lands to enrich the soil.

Since most authors and breeders advocate the enrichment of the soil nutrient status with synthetic fertilizer to facilitate the gain of the improved cassava varieties, and some farming communities relies solely on the natural soil enrichment over the years, this study thus examined the yield and yield related parameters of four newly released cassava varieties at IITA and one local best cassava variety over a period of two planting seasons in a naturally enriched soil by considering the soil nutrient status over two planting seasons.

## MATERIALS AND METHODS

### Experimental location

The experiment was conducted at the Ogbagu Ogume community farms in the 2023 and 2024 early farming seasons in three farm plots: Farms (A, B and Farm C) in the community located at Latitude 5.74737009 and Longitude 6.35451690.

### Source of planting materials

The cassava stems of Hope, Obasanjo2, Baba 70, and Game changer were obtain from the International Institute of Tropical Agriculture (IITA)’s product demonstration exercise for the stability of the newly released cassava varieties in which Ogbagu Ogume was one of the selected communities. One local variety known by the farming community as the local best due to its wide acceptability in the community as the best cassava variety was used as a local check.

### Land preparation, planting and field management

The land vegetation were cleared, trees were cut down and both cleared grasses and trees were allowed to dry properly for one month with the aid of natural sunlight. After which the farms were set on fire to burn the grasses. The burnt fallen trees were packed and kept at the farm boundaries. Selected lands measures 20 m^2^. The cassava stems of the four (4) newly released varieties and one local best variety that are longer than 20 cm were cut into 20 cm sizes each and planted 0.5 cm apart. The farming community practices shifting cultivation by leaving a cultivated land area for at least five (5) years to replenish its nutrient contents, hence no artificial or organic fertilizer is applied to the farms, while irrigation is natural rain fed. The fields were cleared (weeded) of emerging grasses bi monthly till harvest. At dry season, the boundaries of the farms were cleared of any grass or debris to prevent fire encroachment to the farms.

### Experimental design

The experiment was a three (3) way factorial (5 (varieties) x 3 (farm locations) x 2 (farming seasons)) laid in a Randomized Complete Block Design (RCBD) with three replications.

### Pre and post soil analysis

After land preparation, prior to planting for both 2023 and 2024 planting seasons, 14 cm depth soil samples were taken from 10 marked points of the farm covering the entire farm divided into sections. The soils were mixed together to form a composite samples in three replications and sent to the soil analytical laboratory of the Department of Soil Resources Management, Faculty of Agriculture and Forestry, University of Ibadan for the estimation of the soil physico-chemical parameters.

### Data collection and statistical analysis

The soil physico-chemical parameters comprises of soil pH, organic carbon, Nitrogen, Phosphorus, exchangeable acidity, Calcium, Magnesium, Sodium, sand, silt, clay, Manganese, Iron, Copper, and Zinc contents. Data were also taken on the number of stem tubers, stem height and number of stem per stock, the fresh tuber yield in Kg at harvest and extrapolated to hectares following the formular below;

Yield per ha was obtained by converting the yield per plot to hectare below:

1 ha = 10000 m^2^,

20 m^2^ = 20/10000 m^2^ = 0.002 ha,

Hence, yield per 20 m^2^ x 500 = 1 ha equivalence.

Data collected were analysed using Analysis of Variance (ANOVA) and differences in treatment means were separated using Least Significant Differences at 5% level of significance.

## RESULTS

### Growth and yield related performances

Results obtained on the varietal growth and yield related parameters showed that average plant height per stand was recorded in the local best (2.3±0.03) cm, and it was significantly higher than the rest newly released varieties (Table 1). The local best variety also had the highest number of stem per stand (4.07±0.14) which was significantly higher than the rest varieties. Obasanjo2 had the heaviest fresh weight (40.96±0.52) t/ha which was significantly heavier than the rest varieties as the lowest weight was recorded in the local best (19.22±0.52) t/ha. However, average number of roots weighed in the local best (12.47±0.28) was significantly higher than the newly released varieties. As Hope had the lowest number of root tubers per stand (8.4±0.28).

**Table 1.** Growth and yield performances of newly released cassava varieties in Ogume farming community.

| Varieties | Fresh weight (t/ha) | Number of roots | Average stem height | Number of stem per stand |
| --- | --- | --- | --- | --- |
| Baba70 | 33.46b | 9.37b | 2.02b | 3.27bc |
| Game Changer | 34.76b | 9.37b | 1.89c | 3.37b |
| Hope | 34.85b | 8.40c | 1.51d | 2.43d |
| Obasanjo2 | 40.96a | 9.90b | 1.5d | 3.00c |
| Local Best | 19.22c | 12.47a | 2.3a | 4.07a |
| LSD(0.05) | 1.44 | 0.79 | 0.10 | 0.39 |
| SE | 0.52 | 0.28 | 0.03 | 0.14 |
| Variety*Year | 133.50** | 13.36** | 0.12* | 1.08ns |
| Variety*Farms | 7.93ns | 7.22** | 0.05ns | 0.19ns |
| Farms*Year | 28.57* | 3.09ns | 0.11ns | 0.65ns |
| Variety*Year*Farms | 20.94* | 3.94ns | 0.08* | 0.59ns |

Means with same alphabet down the group are not significantly different from each other. LSD: Least Significant Differences, SE: Standard error. Year\*Farms: Interaction between year and farm, Year\*Stage: Interaction between year and stage of analysis. Farms\*Stage: Interaction between farms and stage of analysis.

The interaction between the cultivated varieties and year of cultivation was significant for the average stem height, fresh weight (t/ha) and number of root tubers. The interaction between the varieties and farms cultivated was significant for the number of root tubers harvested. The interaction between the farms cultivated and the year of cultivation was significant for the fresh tuber weight harvested, while the interaction between the varieties cultivated, year of cultivation and the farms cultivated was significant for the fresh tuber weight (t/ha) harvested.

Table 2 showed that the average height of the cassava stems harvested, number of stems per stand , and the number of root tubers harvested were not significantly different across the farms cultivated (farms A, B and C). The fresh weight (t/ha) of the cassava harvested from the Farm A (34.13±0.40) was significantly higher than the rest two farms (farm B and C).

**Table 2.** Effect of farm location and year of planting on the growth and yield performance of newly released cassava varieties.

| Farms | Fresh weight (t/ha) | Number of roots | Average height | Number of stem per stand |
| --- | --- | --- | --- | --- |
| Farm A | 34.13a | 9.72a | 1.83a | 3.24a |
| Farm B | 31.72b | 9.90a | 1.83a | 3.22a |
| Farm C | 32.11b | 10.08a | 1.85a | 3.20a |
| LSD(0.05) | 1.12 | 0.61 | 0.08 | 0.30 |
| SE | 0.40 | 0.22 | 0.03 | 0.11 |
| Year |  |  |  |  |
| One | 34.95a | 9.83a | 1.81a | 3.27a |
| Two | 30.35b | 9.97a | 1.87a | 3.17a |
| LSD(0.05) | 0.91 | 0.5 | 0.07 | 0.24 |
| SE | 0.33 | 0.18 | 0.02 | 0.09 |

Means with same alphabet down the group are not significantly different from each other. LSD: Least Significant Differences, SE: Standard error.

### Soil condition of the cultivated farms across two planting seasons

Table 3 showed that the soil pH, organic carbon, Nitrogen, Phosphorus, exchangeable acidity, Potassium and Sodium contents of the farms cultivated were not significantly different across the two years of cultivation. However, the Calcium content of the soil in the first year (2.03±0.11) was significantly higher than the second year (1.49±0.11) of cultivation. Also, the Magnesium content of the soil in the first year (1.05±0.07) was significantly higher than the second year of cultivation (0.79±0.07). Also, the soil pH, organic carbon, Nitrogen, Phosphorus, exchangeable acidity, Calcium, Magnesium and Sodium contents of the farms were not significantly different across the farms, while the Potassium content recorded in Farm B (0.42±0.04) was statistically similar to Farm C (0.34±0.04) but was significantly higher than the Farm A (0.29±0.04).

**Table 3.**
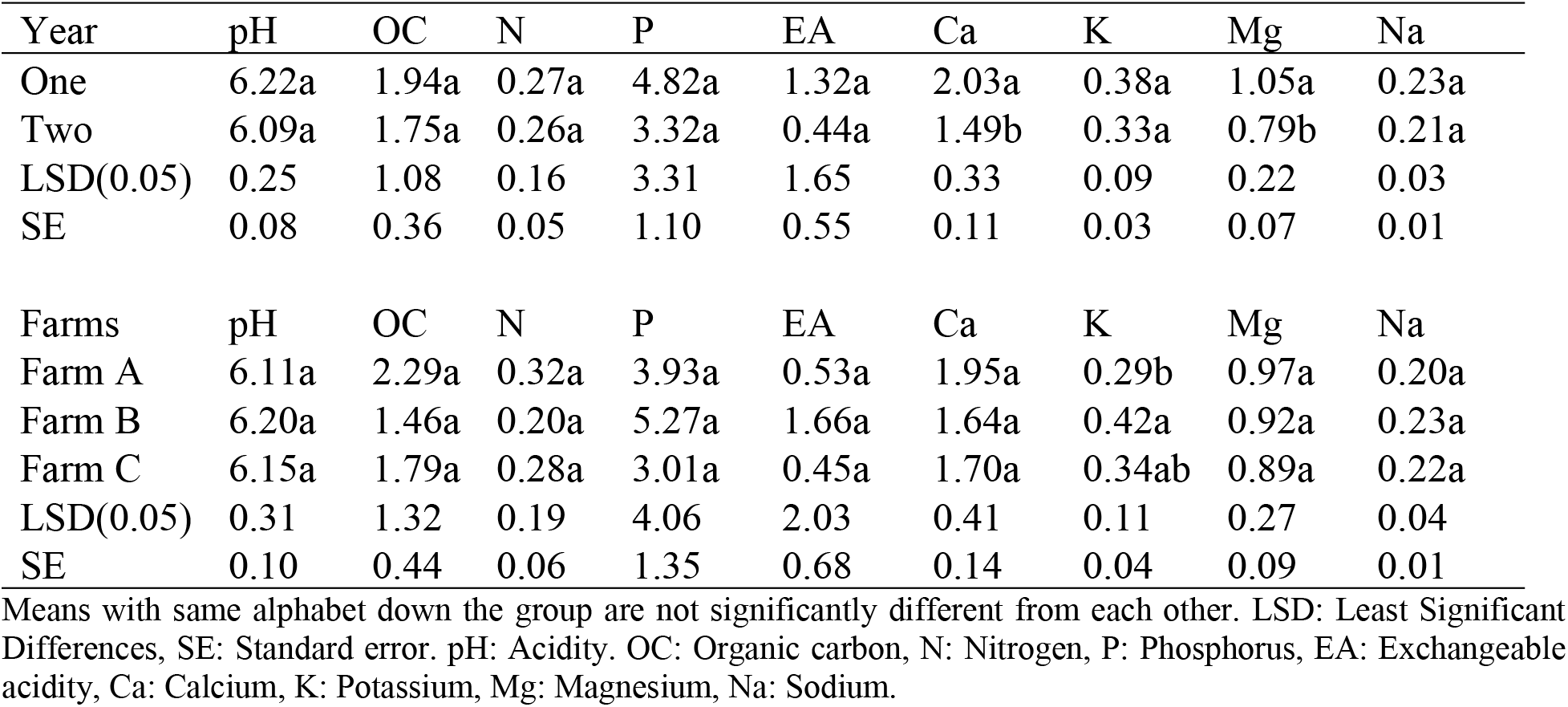
Effect of cassava farm location and year of evaluation on the pH, organic carbon, exchangeable acidity and macronutrient compositions in Ogume farming community.

| Year | pH | OC | N | P | EA | Ca | K | Mg | Na |
| --- | --- | --- | --- | --- | --- | --- | --- | --- | --- |
| One | 6.22a | 1.94a | 0.27a | 4.82a | 1.32a | 2.03a | 0.38a | 1.05a | 0.23a |
| Two | 6.09a | 1.75a | 0.26a | 3.32a | 0.44a | 1.49b | 0.33a | 0.79b | 0.21a |
| LSD(0.05) | 0.25 | 1.08 | 0.16 | 3.31 | 1.65 | 0.33 | 0.09 | 0.22 | 0.03 |
| SE | 0.08 | 0.36 | 0.05 | 1.10 | 0.55 | 0.11 | 0.03 | 0.07 | 0.01 |
| Farms | pH | OC | N | P | EA | Ca | K | Mg | Na |
| Farm A | 6.11a | 2.29a | 0.32a | 3.93a | 0.53a | 1.95a | 0.29b | 0.97a | 0.20a |
| Farm B | 6.20a | 1.46a | 0.20a | 5.27a | 1.66a | 1.64a | 0.42a | 0.92a | 0.23a |
| Farm C | 6.15a | 1.79a | 0.28a | 3.01a | 0.45a | 1.70a | 0.34ab | 0.89a | 0.22a |
| LSD(0.05) | 0.31 | 1.32 | 0.19 | 4.06 | 2.03 | 0.41 | 0.11 | 0.27 | 0.04 |
| SE | 0.10 | 0.44 | 0.06 | 1.35 | 0.68 | 0.14 | 0.04 | 0.09 | 0.01 |

Means with same alphabet down the group are not significantly different from each other. LSD: Least Significant Differences, SE: Standard error. pH: Acidity. OC: Organic carbon, N: Nitrogen, P: Phosphorus, EA: Exchangeable acidity, Ca: Calcium, K: Potassium, Mg: Magnesium, Na: Sodium.

Table 4 showed that apart from the exchangeable acidity which was not significantly different across the pre and post soil stage, the pre soil contents for pH (6.45±0.08), organic carbon (2.56±0.36), Nitrogen (0.41±0.05), Phosphorus (6.15±1.10), Calcium (2.30±0.11), Potassium (0.46±0.03), Magnesium (1.42±0.07), and Sodium (0.27±0.01) was significantly higher than the post soil statuses. The interaction between the year of cultivation and the stage of the soil (pre or post) was significant for the Sodium content.

**Table 4.** Pre and post pH, organic carbon, exchangeable acidity and macronutrient compositions of cassava farms cultivated in Ogume farming community.

| Stage | pH | OC | N | P | EA | Ca | K | Mg | Na |
| --- | --- | --- | --- | --- | --- | --- | --- | --- | --- |
| Pre | 6.45a | 2.65a | 0.41a | 6.15a | 0.51a | 2.30a | 0.46a | 1.42a | 0.27a |
| Post | 5.86b | 1.04b | 0.12b | 1.99b | 1.25a | 0.93b | 0.25b | 0.43b | 0.16b |
| LSD(0.05) | 0.25 | 1.08 | 0.16 | 3.31 | 1.65 | 0.33 | 0.09 | 0.22 | 0.03 |
| SE | 0.08 | 0.36 | 0.05 | 1.10 | 0.55 | 0.11 | 0.03 | 0.07 | 0.01 |
| Interactions |  |  |  |  |  |  |  |  |  |
| Year*Farms | 0.39ns | 3.34ns | 0.05ns | 6.88ns | 3.39ns | 0.17ns | 0.01ns | 0.02ns | 0.01ns |
| Year*Stage | 0.07ns | 2.14ns | 0.04ns | 17.59ns | 4.08ns | 0.01ns | 0.01ns | 0.01ns | 0.01* |
| Farms*Stage | 0.04ns | 0.43ns | 0.01ns | 13.59ns | 3.52ns | 0.18ns | 0.03ns | 0.03ns | 0.01ns |
| Y*F*S | 0.03ns | 2.43ns | 0.04ns | 1.02ns | 3.39ns | 0.25ns | 0.01ns | 0.02ns | 0.01ns |

Means with same alphabet down the group are not significantly different from each other. LSD: Least Significant Differences, SE: Standard error. Year\*Farms: Interaction between year and farm, Year\*Stage: Interaction between year and stage of analysis. Farms\*Stage: Interaction between farms and stage of analysis. pH: Acidity. OC: Organic carbon, N: Nitrogen, P: Phosphorus, EA: Exchangeable acidity, Ca: Calcium, K: Potassium, Mg: Magnesium, Na: Sodium.

Table 5 showed that the sand and silt structures, Manganese, and Copper contents of the cultivated soils was not significant across the years of cultivation. However, the clay content (11.60±0.51) of the second year was significantly higher than the first year (9.23±0.51), Iron (125±2.96) and Zinc (1.93±0.03) contents of the first year of cultivation was also significantly higher than the second year contents with, 112.50±2.96, and 1.82±0.03, respectively. In terms of the three different farm locations cultivated, the sand, silt, Manganese, Iron, Copper, and Zinc contents were not significantly different amongst them, however, the clay content of Farm C (11.50±0.62) was significantly higher than Farm B (9.23±0.62), but was statistically similar to Farm A (10.13±0.62).

**Table 5.** The soil physical structure and microelement compositions of three cassava farm locations in Ogume farming community for two planting seasons.

| Year | Sand | Silt | Clay | Mn | Fe | Cu | Zinc |
| --- | --- | --- | --- | --- | --- | --- | --- |
| One | 79.61a | 10.98a | 9.23b | 64.83a | 125.92a | 1.16a | 1.93a |
| Two | 78.60a | 9.38a | 11.60a | 70.42a | 112.50b | 1.18a | 1.82b |
| LSD(0.05) | 2.54 | 2.01 | 1.52 | 7.89 | 8.88 | 0.13 | 0.09 |
| SE | 0.85 | 0.67 | 0.51 | 2.63 | 2.96 | 0.04 | 0.03 |
| Farms |  |  |  |  |  |  |  |
| Farm A | 80.28a | 9.60a | 10.13ab | 67.25a | 114.50a | 1.19a | 1.91a |
| Farm B | 79.15a | 10.35a | 9.23b | 69.88a | 123.00a | 1.23a | 1.85a |
| Farm C | 77.90a | 10.60a | 11.50a | 65.75a | 120.13a | 1.09a | 1.87a |
| LSD(0.05) | 3.11 | 2.47 | 1.86 | 9.66 | 10.87 | 0.16 | 0.12 |
| SE | 1.04 | 0.82 | 0.62 | 3.22 | 3.62 | 0.05 | 0.04 |

Means with same alphabet down the group are not significantly different from each other. LSD: Least Significant Differences, SE: Standard error. Mn: Manganese, Fe: Iron, Cu: Copper.

The silt (11.25±0.67), Copper (1.63±0.04), and Zinc (2.16±0.03) contents of the pre soil status was significantly higher than the post status with 9.12±0.67 (silt), 0.71±0.04 (Copper) and 1.59±0.03 (Zinc) contents of the post soil statuses (Table 6). However, the Clay (11.92±0.51) and Manganese (73.58±2.63) contents of the post soil status was significantly higher than the pre soil status with 8.92±0.51 and 61.67±2.63 for clay and Manganese, respectively. The interaction between the year of cultivation, farms cultivated and the stage of soil analysis was only significant for the Zinc content.

**Table 6.**
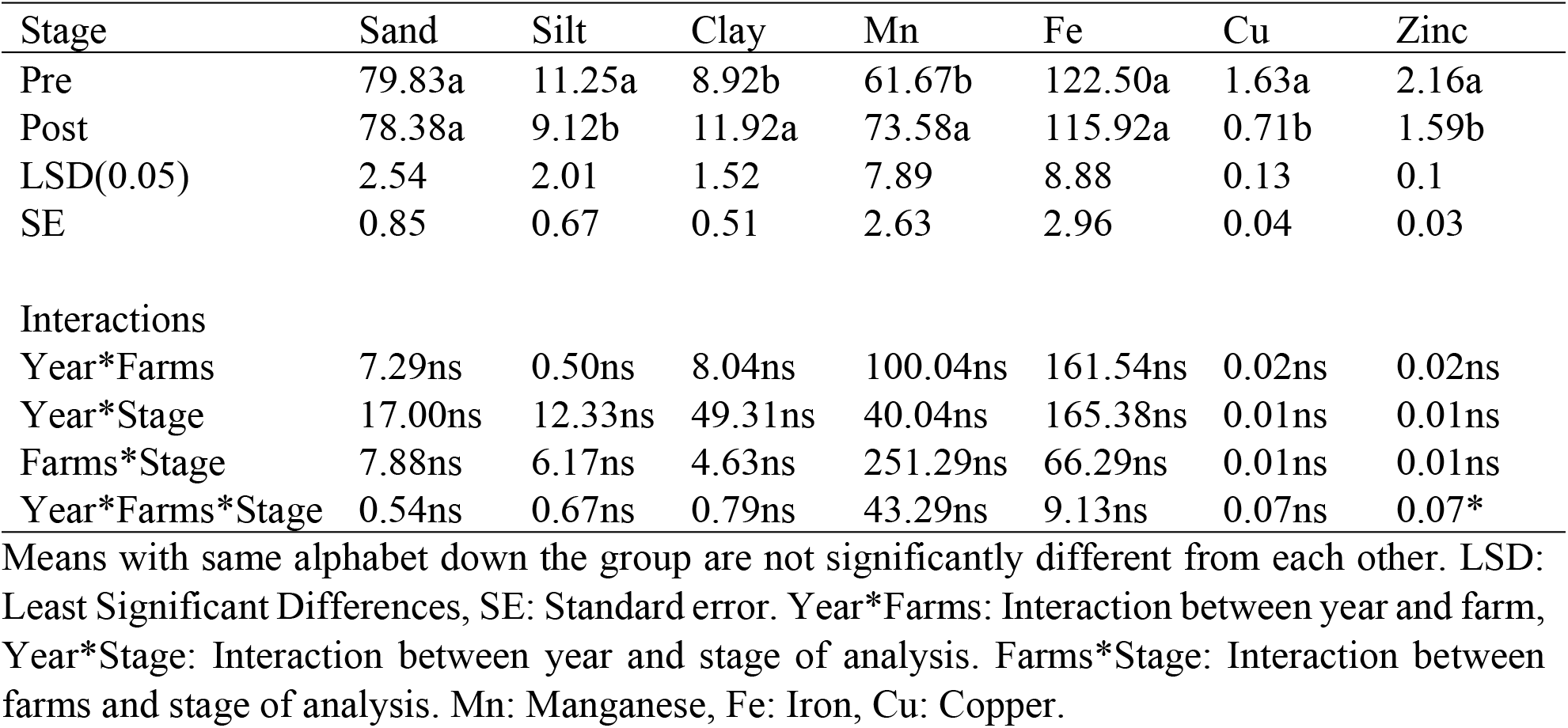
Pre and post soil structures and micronutrient compositions of cassava farms cultivated in Ogume farming community.

## DISCUSSION

Cassava has gained so much importance globally due to the direct utilization of the stem tubers which can be peeled off the back and processed into garri, fufu, tapioca, bread, starch, and ethanol, while the peels can be used as feed for animals and organic composts. The various uses calls for increased cassava output in form of the stem tubers, and given optimum conditions, differences in variety output can be genotypic. Despite Africa being the largest producer of cassava worldwide, the average yield of around 10 t/ha is too low compared to the 26 t/ha obtainable in India (Asia) (15). In this study, the four (4) improved cassava varieties developed by IITA doubled the yield of the local best variety which the local farmers have been cultivating for years and some farmers have shown recalcitrant in adopting and replacing the old variety with the improved ones. An improvement in the yield and yield related parameters of eleven improved cassava varieties developed by the IITA cassava breeding unit relative to local varieties have been earlier reported by Dimkpa et al. (16). One of the major goal of the IITA breeding programs was to identify several constraints militating against crop production in sub-Saharan Africa and develop an improved varieties that are tolerant or resistant to the factors in order to improve the crop yield (17).

In the output recorded, the increased yield was not based on the production of higher number of stem tubers, but on the bulking size of the tubers as seen in the result obtained where the number of the tubers produced per stand in the local variety was higher than the ones produced by the improved varieties. One of the major constraint to the adoption of the improved varieties is the rapid multiplication of the planting material e.g stem. In this regard, the popular stem bundle package in sale is usually 1 m by 12 pieces, and going by this calculation, more income will be generated by the sale of the local variety, however, factoring the double value of yield obtainable by cultivating the improved varieties, the little difference in bundle size is compensated for. This also means that in terms of propagation, the local varieties tend to spread faster as its propagation ratio will be higher than the improved varieties as propagation is based on stem height. This is an area that requires the attention of the breeding institutes to factor in stem sizes in the breeding and selection program as it will help facilitate the cassava seed system in the spread to end users and improve the low adoption rate of improved cassava varieties (18). As at 2011, the adoption rate of improved cassava varieties by farmers in Nigeria stood at 82.2% (19) which decreased to 61% in 2015 (20). Improving the adoption rate is important as the low adoption of improved cassava varieties has been identified as one of the major reasons for the low agricultural production leading to increased poverty level in Nigeria (21). Abdoulaye et al. (5) had also demonstrated that the cassava yield of farming communities that adopted improved cassava varieties was higher than the communities that did not and they linked awareness to the adoption success.

Improved cassava yield is not only a factor of gene effect (superior genotype), but a combination of genotypic and environment. Although, cassava is known to produce appreciably even in a marginal soil, but will ultimately thrive and give better produce in a balanced soil with optimum nutrient conditions. This study considered the performances of the cassava varieties in three farm locations that were fallowed for a period of at least five (5) years to enable proper nutrient replenishment. Farm A stood out above other two farms (B and C) in the yield and yield related parameters of the cassava varieties, while the years of cultivation was only significant in the fresh tuber weight harvested. This shows that the cassava varieties benefited from the abundance of the soil nutrient elements in the first year with small depreciation caused by the plants nutrient uptake, hence the better performance in the first year relative to the second planting season. Although cassava gives appreciable yield in marginal soil (22), the output is far better in a rich soil especially soil with good amount of N, P, and K, the reason why farmers constantly resort to the application of NPK inorganic fertilizer (23). However, these elements were not lacking in this farming community due to the shifting cultivation farming system which aided the replenishment of the nutrients.

In the two planting seasons, the macro elements reduced in content as a result of replanting. The depreciation in the macro elements was further substantiated in the reduction in all the elements relative to the post status of the soil. This goes along with the feeding habit of cassava which is a deep feeder, and the amount of nutrient uptake is more than the decomposition replenishment with the one year period. On the microelements, there were no significant changes in the contents of Mn and Cu in both years of cultivation, however, while considering the pre and post soil nutrient contents the Mn content of the post was higher than the pre status, which explains why the value improved in the second year which could be that decomposing cassava leaves has the ability to release Mn to the soil. The reduction in the macronutrient content of the soil in the second year is as expected as the plants take up the elements in large quantity from the soil (22). Also, the lower nutrient content (majority of the elements) of the second season explains why the farming community only farms in a location once and abandons the site for the next 5 years before they return to it as they do not apply inorganic fertilizer to compensate low nutrient status caused by cassava uptake (24). Despite the success recorded in this practice, it is only feasible for farming communities with extensive arable land areas, as those with limited land areas will resort to inorganic fertilization or an integrated nutrient management system, and NPK fertilizers has been greatly advocated due to the large amount of K taken up by cassava stems during bulking (24; 14).

The sand, silt and clay percentage obtained for both two planting seasons showed that the soil type belong to the Loamy Sand textural class, which explains why the soil has adequate organic matter content for crop production, and this emanated from the fallow system operated by the farming community, allowing the decomposition of organic matters and the process of debris burning releases further nutrient to the soil in ash form. Abass et al. (25) has stated that cassava requires a loose-textured soil to enhance the penetration of the roots and its bulking, and these were the soil type for this study which indicates that differences in the genotypic performance was more of varietal differences.

## CONCLUSION

The breeding program of research institutes like IITA has yielded fruit in the development of improved cassava varieties that addressed yield gaps in Nigeria as the yield obtained from the improved varieties were higher than the local best variety. However, the success gained could be hampered by low adoption rate due to low propagation ratio (short stem sizes), awareness campaign robustness, and further educating the farmers on the importance of good soil nutrient status to achieve optimum yield. It is thereby recommended that stem size should be factored into further improving these high yielding varieties for ease of dissemination and there should be more robust collaboration with extension agents in the dissemination processes.

## Author Contributions

Conceptualization: O.C.O., O.S.A., T.O.O., A.S.C., A.S.I and M.E.D.-O. Data Curation: O.C.O. O.S.A. and A.S.C. Methodology: O.C.O., O.S.A., T.O.O., A.S.C., A.S.I and M.E.D.-O.

## Formal Analysis

O.C.O. Writing Original Draft: O.C.O. Writing—Review and Editing: O.C.O., O.S.A., T.O.O., A.S.C., A.S.I and M.E.D.-O. All authors have read and agreed to the published version of the manuscript.

## Institutional Review Board Statement

Not applicable. Informed Consent Statement: Not applicable.

## Data Availability Statement

All data used during the study are available from the author Ossai, C. O. by request.

## Acknowledgments

This study was designed for the evaluation of newly released cassava varieties under the Building an Economically Integrated Sustainable Cassava Seed System (BASICS) in Nigeria funded by the Bill and Melinda Gates Foundation. Thus, we acknowledged BMGF for the BASICS Project award to IITA.

